# A habenula-enriched GPCR, GPR151, regulates behavioral sensitivity to inflammation

**DOI:** 10.64898/2026.08.02.742327

**Authors:** Lupita Rios, You-Hsin Lin, Laura Yuan, Yash Sharma, Hailey Arias, Fizza Jeddy, Sravya Thotakura, Aiden Geleta, Raghav Rajesh, Steven Shabel

## Abstract

**Background:** Inflammation-associated depression is a subtype of major depressive disorder that is often resistant to conventional pharmacotherapies, which act in a regionally non-specific manner and therefore also produce unwanted side effects. Here we test GPR151, an orphan GPCR associated with inflammation and highly expressed in the habenula—a region linked to negative valence and depression—as a therapeutic target for inflammation-associated depression.

**Methods:** We integrated mouse and human habenular expression analyses with genetic loss-of-function and adult habenular re-expression approaches in mice. Gpr151 knockout mice and littermate controls were exposed to lipopolysaccharide (LPS) inflammatory challenge and assessed for stress coping and motivated behavior, body weight loss, and peripheral immune activation. To test whether adult habenular GPR151 expression is sufficient to restore inflammation-associated behavioral vulnerability, GPR151 was re-expressed in the habenula of knockout mice.

**Results:** GPR151 was exceptionally enriched in the habenula and showed conserved topographic organization and similar expression relationships with habenular marker genes in mice and humans. Following LPS challenge, male Gpr151 knockout mice showed reduced passive coping despite body weight loss and immune activation comparable to littermate controls. Adult habenular GPR151 re-expression increased LPS-induced amotivation in male knockout mice without increasing LPS-induced weight loss or immune activation. Female Gpr151 knockout mice also showed reduced passive coping after LPS challenge; however, habenular GPR151 re-expression was insufficient to increase LPS-induced amotivation in females.

**Conclusions:** These findings identify GPR151 as a conserved, regionally enriched regulator of behavioral sensitivity to inflammatory challenge and support GPR151 as a candidate therapeutic target for inflammation-associated depression.

## Introduction

Major depressive disorder is a leading cause of disability worldwide^1^, yet is mechanistically and clinically heterogeneous^2,3^. A substantial subset of patients exhibits peripheral inflammation which has been linked to greater somatic symptoms and poorer treatment response^4–6^. Meta-analytic work suggests that roughly a quarter of depressed patients meet criteria for low-grade inflammation, with an even larger fraction showing at least mildly elevated inflammation, consistent with inflammation being relevant for a meaningful subgroup rather than a rare comorbidity^7^. Additionally, converging evidence indicates that inducing strong, systemic inflammation increases the risk of depressive symptoms^8–10^, and inflammatory challenge in non-human animals induces amotivation, a core symptom of depression^11–13^. Collectively, these observations support a stratified view of depression in which inflammatory state is a tractable axis for patient segmentation and potentially for precise therapeutics.

In depressed patients, inflammation is associated with altered function in networks that subserve reward, motivation, and goal-directed behavior^14–18^. These observations point to the potential value of interventions that directly modulate brain regions governing negative valence and motivational state. Pharmacotherapies that act specifically on these regions may be more effective and have fewer side effects than currently available treatments.

One such brain region is the habenula, a diencephalic structure that has emerged as a key regulator of negative valence, non-reward processing, and motivation through its control of monoaminergic brain regions^19–22^. Consistent with this view, human and preclinical studies implicate habenular dysfunction in amotivation and depression^20,23–50^. The habenula’s strong influence over neuromodulatory systems makes it an attractive locus for region-selective interventions aimed at negative affect and amotivation. Furthermore, although the habenula has not been studied extensively in neuroimmune regulation, several reports indicate that inflammatory cytokine signaling within the medial habenula (MHb) or LHb is altered by stress, inflammatory challenge, and drug withdrawal, and can contribute to depression-relevant behavioral phenotypes^45,46,51–55^.

G protein-coupled receptors (GPCRs) represent an unusually “druggable” protein class and therefore a pragmatic entry point for developing brain-region-biased therapeutics^56^. Within this class, receptors with highly restricted neuroanatomical expression patterns are particularly appealing because they raise the possibility of modulating specific brain regions while minimizing off-target effects. GPR151 is an orphan GPCR that stands out for its strong habenular expression in the brain^55,57–59^, with additional expression reported in sensory ganglia and spinal cord^59^, where it can modulate nociception and inflammation^60–62^. Human genetic data also provide suggestive support for a link between *GPR151* and depression: in a large depression GWAS meta-analysis, *GPR151* showed a gene-based association^63^ (although it did not meet the genome-wide, multiple-testing statistical threshold). However, whether GPR151 contributes to inflammation-associated motivational deficits—and whether it does so through adult habenular mechanisms that would be therapeutically addressable—is unclear.

Here we test the hypothesis that GPR151 is a habenula-enriched mediator of inflammation-associated, depression-relevant behavior. Using lipopolysaccharide (LPS) as an established inflammatory challenge^64,65^, we first assessed the behavioral consequences of genetic loss of GPR151 and separated motivational phenotypes from sickness by measuring LPS-induced weight loss and immune activation alongside behavioral assays. We then evaluated adult, region-specific sufficiency by restoring GPR151 expression selectively within the habenula and measuring LPS-induced amotivation across complementary behavioral domains, including stress coping, exploratory locomotion, and reward sensitivity in both sexes. This design addresses a translationally central question: can modulation of a habenula-enriched orphan GPCR affect behavioral sensitivity to inflammation? Together, these experiments position GPR151 within an emerging mechanistic framework that links peripheral inflammatory state to specific brain regions governing motivated behavior and support a member of a tractable receptor class for therapeutic exploration in inflammation-associated depression.

## Methods and Materials

### Animals

Heterozygous *Gpr151^+/−^* mice^66^ (Jackson Labs) were used for breeding to produce wild-type (WT) and *Gpr151^−/−^* knockout littermates used for experiments. Mice were group-housed (2– 5 per cage) under a 12:12 h light/dark cycle with ad libitum access to food and water until ∼ 1 week before the start of baseline sucrose preference testing, at which point they were singly housed for the remainder of the experiment. All procedures were approved by the Institutional Animal Care and Use Committee at UT Southwestern Medical Center and conformed to NIH guidelines.

Tg(*Gpr151*-cre) mice, generated by BAC transgenesis to express Cre recombinase under the control of the Gpr151 promoter, were kindly provided by Dr. Shigeyoshi Itohara (RIKEN). To eliminate endogenous Gpr151 while maintaining cre expression, Tg(Gpr151-cre) mice were crossed with *Gpr151^−/−^* mice. Offspring carrying both the Cre transgene and Gpr151 null alleles (Cre+/KO) were used for experiments.

One female and one male mouse in the GPR151^Hb^ re-expression groups were excluded from all analyses, except the habenula expression vs behavior scatterplots, due to lack of GPR151 expression in the habenula.

### LPS administration

Lipopolysaccharide (0.8 mg/kg, intraperitoneal; Sigma, Cat. #L3129-10MG) or sterile saline was administered once daily for three consecutive days. In viral cohorts, injections began 3-4 weeks after surgery. Behavioral testing commenced 24 hours after the final injection, a time point chosen based on prior studies demonstrating resolution of acute sickness behaviors within this window^13^. The selected dose has been validated in previous work to induce robust inflammation and depression-like behavior^67^. A three-day schedule was used because a similar three-day LPS schedule induced *Gpr151* expression in sensory ganglia 24 hours later^68^ and because neuropathic pain manipulations induce *Gpr151* expression in sensory ganglia and/or spinal cord after 3-7 days^60,61,66,69,70^.

### Behavioral testing

All behavioral testing was performed during the light phase. Mice were habituated to the testing room for at least 1 h prior to experiments. Order of testing was counterbalanced across groups. Experimenters were blinded to genotype and treatment during behavioral scoring and analysis.

#### Open Field Test (OFT)

Locomotor activity and center time were assessed in a square open-field arena (40 × 40 cm) under consistent lighting. Each mouse was placed in the center and allowed to explore for 10 min. Total distance traveled and time spent in the center versus periphery were recorded using EthoVision XT software (Noldus Information Technology). The arena was cleaned with antiseptic solution between trials.

#### Forced Swim Test (FST)

Mice were placed individually in a transparent cylindrical tank (20 cm diameter, 25 cm height) filled with water (23–25 °C) to a depth of 15 cm. Each session lasted 6 min, and immobility during the final 4 min was scored by an observer blinded to genotype and treatment. Water was replaced between trials. Mice were tested in the FST 2-3 hours after OFT testing. One WT male and three male GPR151^Hb^ mice given LPS were not run in the FST due to extreme lethargy.

#### Sucrose Preference Test (SPT)

Mice were first habituated to two water bottles for two days, followed by two days with continuous access to bottles containing 1% sucrose solution. Baseline preference testing began on Day 1 with one water bottle and one sucrose bottle. Testing continued for 5 consecutive days, with bottles weighed daily and positions counterbalanced. Five days after the start of sucrose preference testing, LPS/saline injections commenced, and sucrose preference was recorded throughout injections and subsequent behavioral assays. Although all mice were tested for sucrose preference, only data from the last two cohorts of each type of experiment (KO/WT and viral re-expression) were analyzed due to malfunctioning sipper bottles in previous cohorts.

Please see Supplementary Methods and Materials for more information.

## Results

### Cross-species conservation of habenular *GPR151* expression supports its translational potential

To examine *GPR151* expression in the mouse habenula we performed in situ hybridization in male (Fig. 1A) and female mice (Supplementary Fig. 1A). The expression pattern matched that of previous studies^57,59^ and the Allen Brain Atlas^71^: highly enriched expression in the ventral MHb, at the MHb-LHb border, and in scattered cells in the LHb. Although *GPR151* is highly enriched in the mouse^71,72^ and human habenula^55^, compared to the rest of the brain, it is unclear how its enrichment compares to other GPCRs in other brain regions. To examine this, we used the Allen Brain Atlas database^71^ to find the GPCR with the highest enrichment value in every brain region (*N* = 220 regions) and found that *GPR151* expression in the habenula is the most enriched GPCR of any GPCR in any brain region (Fig. 1B). Thus, pharmacotherapies that target GPR151 function may offer an unusual level of anatomical and functional specificity.

**Fig. 1.**
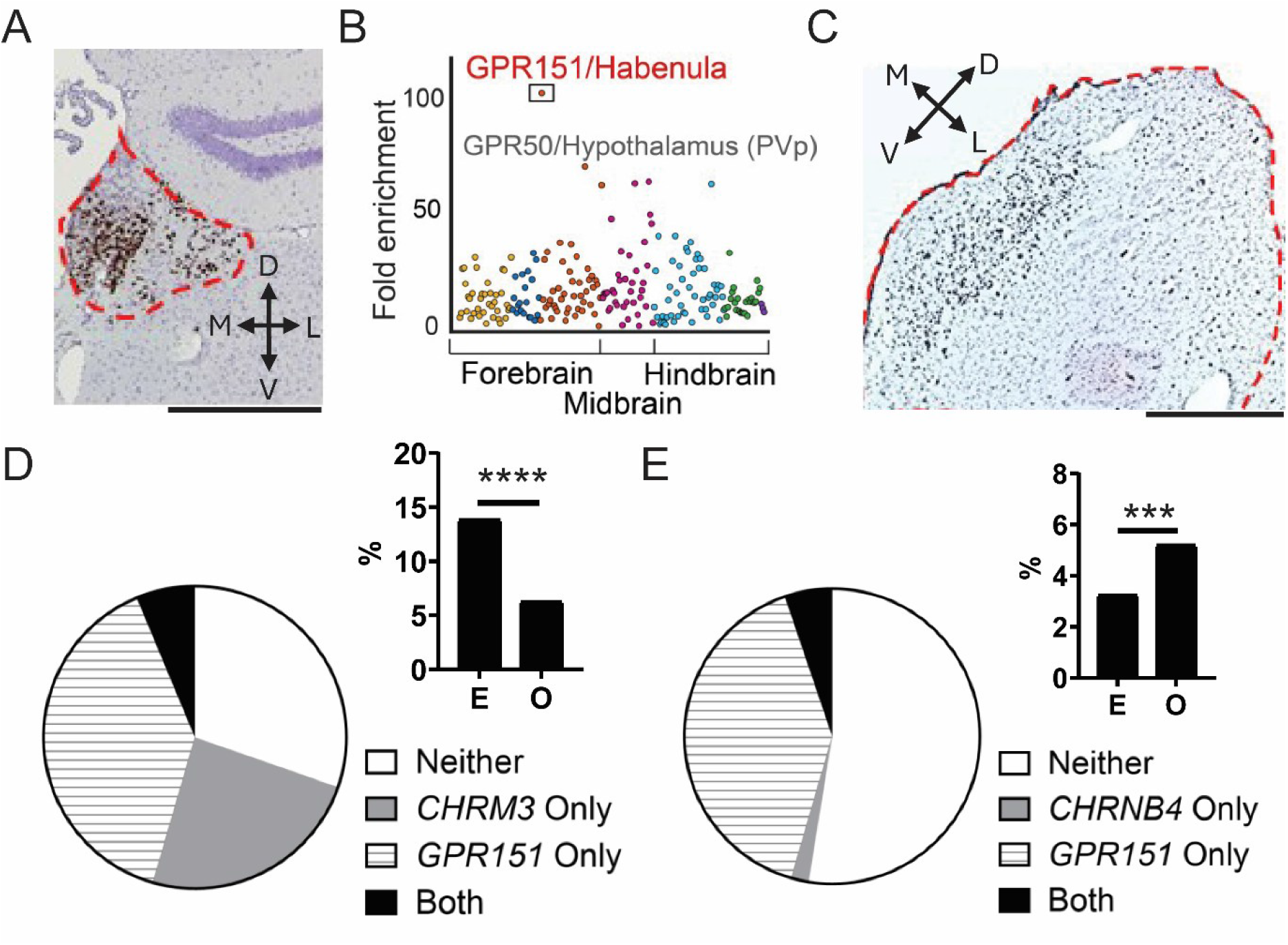
Expression of GPR151 in the mouse and human habenula. **A**, In situ hybridization for GPR151 (brown) in a coronal slice from an adult male mouse. Red dashes outline the habenula. Scale, 0.5 mm. **B**, Enrichment of GPCRs in the mouse brain. Each dot shows the enrichment value of the most enriched GPCR in a brain region compared to the rest of the brain (N = 220 brain regions), according to the Allen Brain Atlas database (see Methods). Colors from left to right represent the cerebral cortex, cerebral nuclei (striatum, pallidum), interbrain (diencephalon), midbrain, hindbrain (except cerebellum), cerebellar cortex, and cerebellar nuclei. PVp, posterior periventricular hypothalamus (GPR50 ranked second), used to help illustrate the analysis. **C**, In situ hybridization for GPR151 (brown) in the human habenula. Red dashes and scale as in A. **D**, Percentage of human habenula neurons expressing GPR151 and/or CHRM3. ‘E’ denotes expected co-expression of the two genes based on the joint probability of their individual expression likelihoods. ‘O’ denotes observed co-expression. ꭕ2 = 87.4, *P* = 8.9 × 10^−21^, N = 2766 neurons. **E**, Percentage of human habenula neurons expressing GPR151 and/or CHRNB4, a marker of MHb cholinergic neurons. ‘E’ and ‘O’, as in D. ꭕ2 = 13.2, *P* = 2.8 × 10^−4^, *N* = 2766. ***, P < .001, ****, P < .0001.

Conserved habenular enrichment and amino acid sequence (∼ 83% identity between rodents and humans^57^) suggest conserved function of GPR151. However, it is unclear if *GPR151* is expressed in similar types of habenular neurons in mice and humans. To determine if *GPR151* is expressed in similar habenular neurons in humans, we performed in situ hybridization for *GPR151* in the human habenula. Similar to its expression in mice^57,71^, *GPR151* was highly expressed in the ventral MHb, at its border with the LHb, and scattered cells in the LHb (Fig. 1C). To further examine the expression of *GPR151* in habenular neurons in humans, we performed single-nucleus RNA sequencing of the human habenula (*N* = 2766 neurons from one human habenula) to determine if the pattern of *GPR151* expression was similar to that of mice. In mice, *GPR151* and *CHRM3*, encoding muscarinic acetylcholine receptor 3, tend to be expressed in different LHb neuronal populations^73^, and *GPR151* expression is high in acetylcholine-releasing neurons in the MHb^57,74,75^. These patterns held in the human habenula: overlap of *GPR151* and *CHRM3* was less than expected by chance (i.e., the actual percentage of neurons that co-expressed *GPR151* and *CHRM3* was less than the joint probability of co-expression given the individual percentages of neurons that expressed *GPR151* or *CHRM3*; Fig. 1D; Supplementary Fig. 1B,C; also true when restricting the analysis to putative LHb neurons; ꭕ2 = 48.2, *P* = 3.9 × 10^−12^, *N* = 2180 neurons) and *GPR151* was co-expressed in cholinergic neurons (using *CHRNB4* as a marker of cholinergic MHb neurons^76,77^; Fig. 1E; Supplementary Fig. 1C; we obtained similar results when using choline acetyltransferase instead of *CHRNB4* as a marker of cholinergic neurons, ꭕ2 = 13.8, *P* = 2.0 × 10^−4^). We also replicated these results using a recently published human habenula snRNA-seq dataset^76^ (Supplementary Fig. 1D-I). Together these data suggest that *GPR151* is expressed in similar neuron types in the human habenula as the mouse habenula, and therefore, is likely to have conserved function in humans.

### *Gpr151^−/−^* mice show reduced passive coping in the forced swim test after LPS treatment

Previous studies indicate a link between *Gpr151* and inflammation^60–62,68^, which is thought to play a causal role in depressive disorders^78^. Additionally, several lines of evidence indicate a role for the habenula in depression and amotivation^19–21^. Therefore, we hypothesized that GPR151 contributes to inflammation-associated depression and amotivation. To test this hypothesis, we used LPS to induce inflammation in *Gpr151* knockout mice and control littermates and measured behavior 24 hours after the final LPS injection: immobility in the forced swim test and locomotion and center time in the open field test (Fig. 2A). We also tested sucrose preference during the entire LPS injection regimen to measure changes in hedonic behavior (Fig. 2A). Consistent with the hypothesis, LPS caused an altered motivational response to stress (i.e., less immobility/passive coping) in male *Gpr151^−/−^* mice (Fig. 2B). There was reduced locomotion after LPS in both groups and slightly more locomotion in male knockout mice, independent of injection (Fig. 2C). Furthermore, there was no interaction between genotype and injection on locomotor activity in males (Fig. 2C) and no negative relationship between locomotion and immobility in the forced swim test (Fig. 2D), indicating that the effect in the forced swim test was not due to a general change in motor activity. There was also no interaction between genotype and injection in center time in the open field and sucrose preference, a measure of hedonic sensitivity, but main effects of injection on both measures (Fig. 2E,F), indicating that LPS affected these behaviors similarly in wild-type and *Gpr151*^−/−^ mice.

**Fig. 2.**
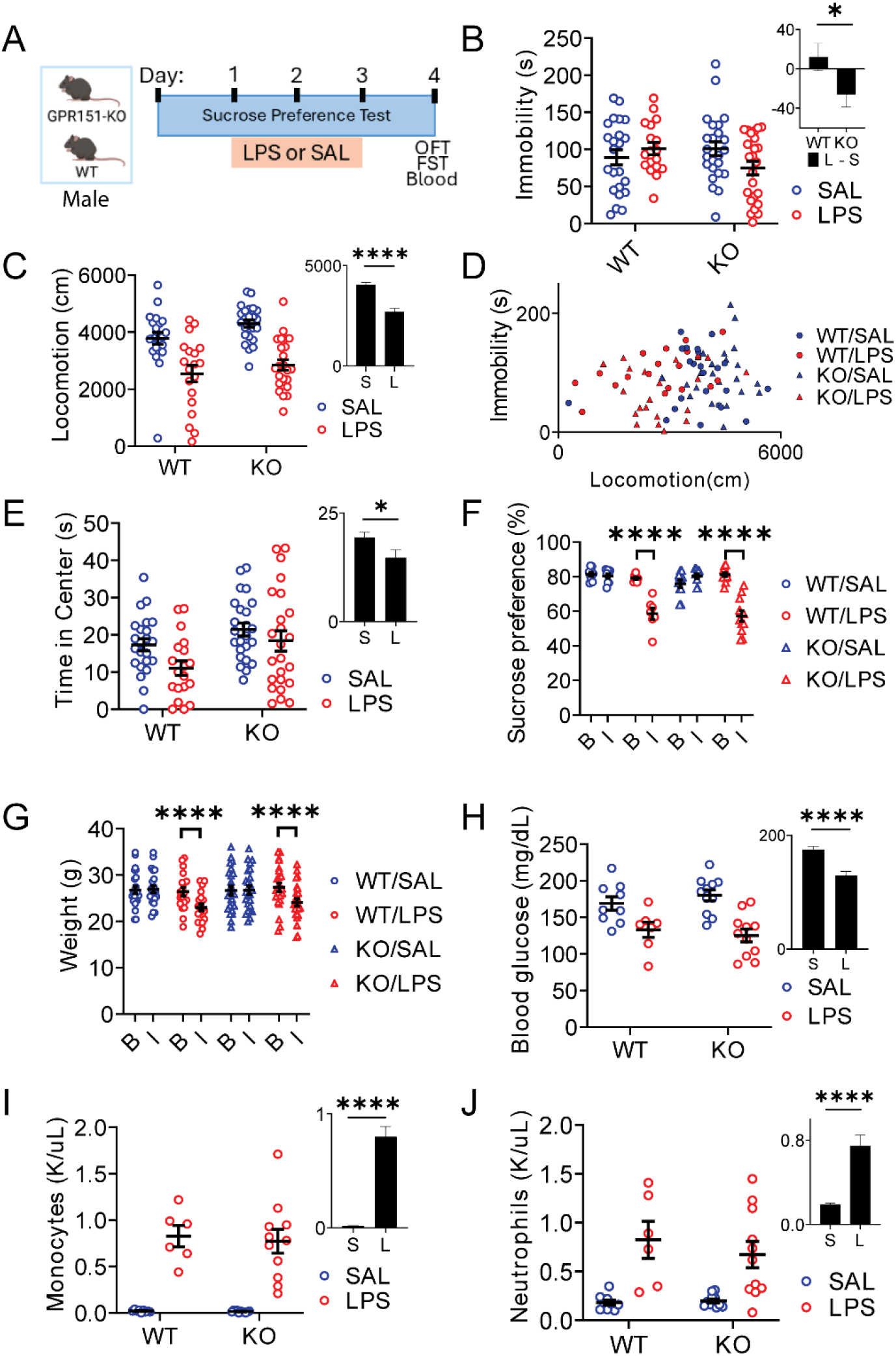
Behavioral and physiological responses to LPS in male *Gpr151* knockout mice. **A**, Experimental procedures. LPS (0.8 mg/kg, i.p.) was administered once per day for three days. Behavior (open field test and forced swim test), blood glucose, and immune markers were measured 24 hours after the last LPS injection. **B**, Immobility in the forced swim test. Inset shows the modeled injection effect size (LPS − Saline), shown separately for WT and KO (estimated marginal mean differences), illustrating the Genotype × Injection interaction. 2-way ANOVA, Genotype x Injection, *P* < .05; *n* = 23 WT/Saline, 18 WT/LPS, 24 KO/Saline, and 24 KO/LPS mice. **C**, Locomotion in the open field test. Inset shows the Saline and LPS estimated marginal means (collapsed across WT/KO), illustrating the main effect of Injection. 2-way ANOVA, Genotype x Injection, *P* > .05, Main effect of Genotype, *P* < .05; *n* = 23 WT/Saline, 19 WT/LPS, 24 KO/Saline, and 24 KO/LPS mice. **D**, Scatterplot of locomotion in the open field test and immobility in the forced swim test. *r* = 0.2, *P* = .07. Mice as in B. **E**, Time in the center of the open field. Inset similar to C. 2-way ANOVA, Genotype x Injection, P > .05. Mice as in C. **F**, Sucrose preference at baseline, ‘B’, and during the 3-day LPS regimen (average of 3 days), ‘I’. 3-way RM ANOVA, Genotype x Injection x Time, P > .05. *n* = 9 WT/Saline, 7 WT/LPS, 11 KO/Saline, 11 KO/LPS mice. **G**, Body weight at baseline and during the 3-day LPS regimen (average of 3 days). 3-way RM ANOVA, Genotype x Injection x Time, P > .05. Mice as in C. **H**, Blood glucose measured 30 minutes after the last behavioral test. Inset similar to C. 2-way ANOVA, Genotype x Injection, P > .05. *n* = 9 WT/Saline, 7 WT/LPS, 11 KO/Saline, 11 KO/LPS mice. **I**, Monocyte levels in blood measured 30 minutes after the last behavioral test. Inset similar to C. 2-way ANOVA, Genotype x Injection, P > .05. *n* = 9 WT/Saline, 6 WT/LPS, 11 KO/Saline, 11 KO/LPS mice. **J**, Neutrophil levels in blood measured 30 minutes after the last behavioral test. Inset similar to C. 2-way ANOVA, Genotype x Injection, P > .05. Mice as in I. Error bars, SEM, in all figures. *, P < .05, **, P < .01, ***, P < .001, ****, P < .0001 in all figures. Please see Supplementary Table 1 for more statistical information. Insets were chosen to show information and effects that could not be shown in the main panels.

The similar effects of LPS on locomotion, center time, and sucrose preference in knockout and wild-type mice suggested that the differential response to LPS in the forced swim test was not due to a general insensitivity to LPS. Consistent with this possibility, male knockout mice had similar decreases in body weight, blood glucose, and immune activation in response to the LPS regimen as WT mice (Fig. 2G-J).

Also consistent with the hypothesis that GPR151 contributes to inflammation-associated amotivation, female *Gpr151^−/−^* mice had a similarly altered response to LPS in the forced swim test as male *Gpr151^−/−^* mice (Fig. 3A,B) that was not due to altered motor activity (Fig. 3C,D). Because both sexes showed an interaction between genotype and injection (saline/LPS) in the forced swim test and 3-way ANOVA found no moderation by sex (3-way ANOVA: Sex x Genotype x Injection, F(1,172) = .001, *P* = .98), we pooled their data and found that *Gpr151^−/−^* mice had less immobility in the forced swim test than wild-type mice in response to LPS (2-way ANOVA; Genotype x Injection, F(1,176) = 7.8, *P* = .006; Post hoc simple-effects analysis: WT/LPS vs KO/LPS, mean difference = 21.1, t(176) = 2.3, *P* = .02).

**Fig. 3.**
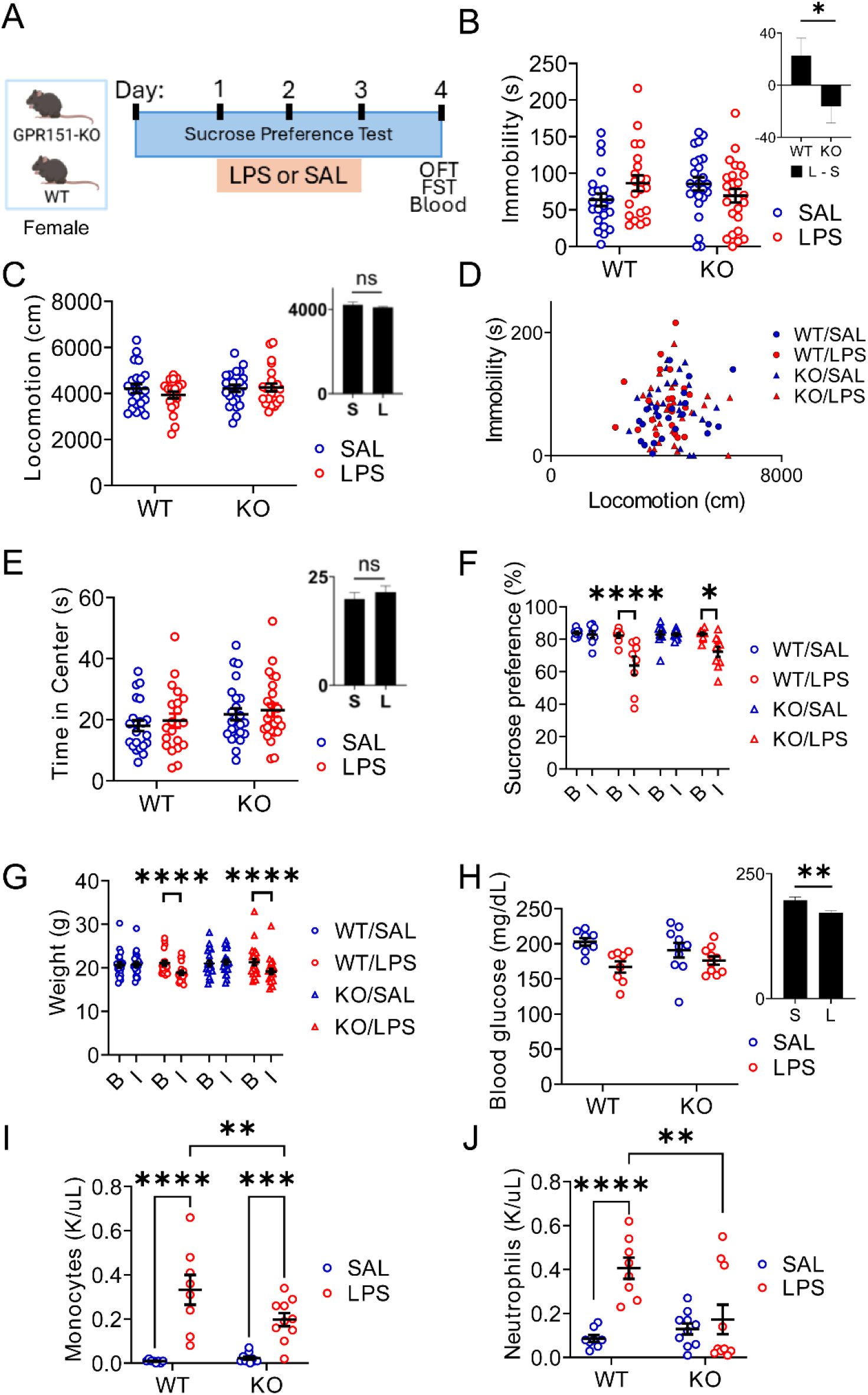
Behavioral and physiological responses to LPS in female *Gpr151* knockout mice. **A**, Experimental procedures, as in Fig. 2. **B**, Immobility in the forced swim test. Inset shows the modeled injection effect size (LPS − Saline), shown separately for WT and KO (estimated marginal mean differences), illustrating the Genotype × Injection interaction. 2-way ANOVA, Genotype x Injection, *P* = .04. *n* = 22 WT/Saline, 21 WT/LPS, 24 KO/Saline, 24 KO/LPS mice. **C**, Locomotion in the open field test. Inset shows the Saline and LPS estimated marginal means (collapsed across WT/KO), illustrating the main effect of Injection. 2-way ANOVA, Genotype x Injection, P > .05. Mice as in B. **D**, Scatterplot of locomotion in the open field test and immobility in the forced swim test. *r* = .06, *P* > .05. Mice as in B. **E**, Time in the center of the open field. Inset similar to C. 2-way ANOVA, Genotype x Injection, P > .05. Mice as in C. **F**, Sucrose preference, as in Fig. 2. 3-way RM ANOVA, Genotype x Injection x Time, P > .05. *n* = 8 WT/Saline, 8 WT/LPS, 10 KO/Saline, and 10 KO/LPS mice. **G**, Body weight, as in Fig. 2. 3-way RM ANOVA, Genotype x Injection x Time, P > .05. Mice as in B. **H**, Blood glucose, as in Fig. 2. Inset similar to C. 2-way ANOVA, Genotype x Injection, P > .05. *n* = 8 WT/Saline, 8 WT/LPS, 10 KO/Saline, 10 KO/LPS mice. **I**, Monocyte levels, as in Fig. 2. 2-way ANOVA, Genotype x Injection, *P* < .05. Mice as in H. **J**, Neutrophil levels, as in Fig. 2. 2-way ANOVA, Genotype x Injection, *P* < .01. Mice as in H. Please see Supplementary Table 1 for more statistical information.

Other measures showed that female mice had an overall reduced response to LPS under these conditions compared to male mice: although they reduced sucrose preference, lost weight, and reduced blood glucose in response to LPS (Fig. 3F-H), they showed no effect of LPS on locomotion nor center time in the open field test (Fig. 3C,E). Females also had less of an increase in monocytes and neutrophils after LPS than males (Fig. 3I,J; Monocytes: Sex x Injection, *P* < .0001; Neutrophils: Sex x Injection, *P* = .002), and notably, female knockout mice showed an even smaller increase in monocytes and neutrophils after LPS than female wild-type mice (Fig. 3I,J).

### Re-expression of GPR151 in the habenula increases inflammation-associated amotivation in male *Gpr151*^−/−^ mice

Differences between wild-type and *Gpr151*^−/−^ mice could be due to developmental differences. To test whether GPR151 expression in the adult habenula affects inflammation-associated amotivation, we re-expressed GPR151 in the adult habenula using *Gpr151*^−/−^ mice that were crossed to *Gpr151*-cre mice (BAC transgenic) and microinfusion of AAVs expressing GPR151 in a cre-dependent manner in the habenula (serotypes 1 and 9 for expression in MHb and LHb; Fig. 4A; “GPR151^Hb^”). This strategy resulted in selective re-expression of GPR151 in the adult habenula and axons within midbrain targets of the habenula (Fig. 4B; e.g., interpeduncular nucleus, IPN, and rostromedial tegmentum, RMTg), as expected^57,79^. Control *Gpr151* knockout/*Gpr151*-cre mice were given habenular microinfusions of the same serotype AAVs expressing a fluorescent reporter (“KO”; Fig. 4A). Consistent with the hypothesis, male GPR151^Hb^ mice had more inflammation-associated amotivation in the forced swim test than male KO mice (Fig. 4C), and GPR151 expression in the habenula was correlated with immobility in LPS-treated mice, but not saline mice (Fig. 4D; Supplementary Fig. 2A,E; LPS: Spearman, *r_s_* = .73, *P* < .001, *N* = 17 mice; Saline: *r_s_* = .28, *P* = .25, *N* = 19 mice). Male GPR151^Hb^ mice were also more affected by LPS than male KO mice when measuring locomotion in the open field test (Fig. 4E,F; Supplementary Fig. 2B,F), but not center time (Fig. 4H), although again, there was no significant relationship between locomotion in the open field test and immobility in the forced swim test (Fig. 4G), indicating that differences in immobility in the forced swim test were not due to general hypoactivity. Male GPR151^Hb^ mice were also more affected by LPS than male KO mice in the sucrose preference test (Fig. 4I) and the difference between groups only emerged after the second LPS injection (Fig. 4K-M). Similar to results from the forced swim test, GPR151 expression in the habenula was strongly correlated with a reduction in sucrose preference in LPS-treated mice (Fig. 4J, Supplementary Fig. 2C,G). There was no difference in body weights (Fig. 4N) and blood glucose levels of KO and GPR151^Hb^ mice after LPS (Fig. 4O), and no increase in the immune response to LPS in GPR151^Hb^ mice compared to KO mice (Fig. 4P,Q; although there was a surprising reduction in monocyte levels in male GPR151^Hb^/LPS mice compared to KO/LPS mice), indicating that the hypersensitivity to LPS in male GPR151^Hb^ in the forced swim, open field (locomotion), and sucrose preference tests was not due to a hypersensitive immune response.

**Fig. 4.**
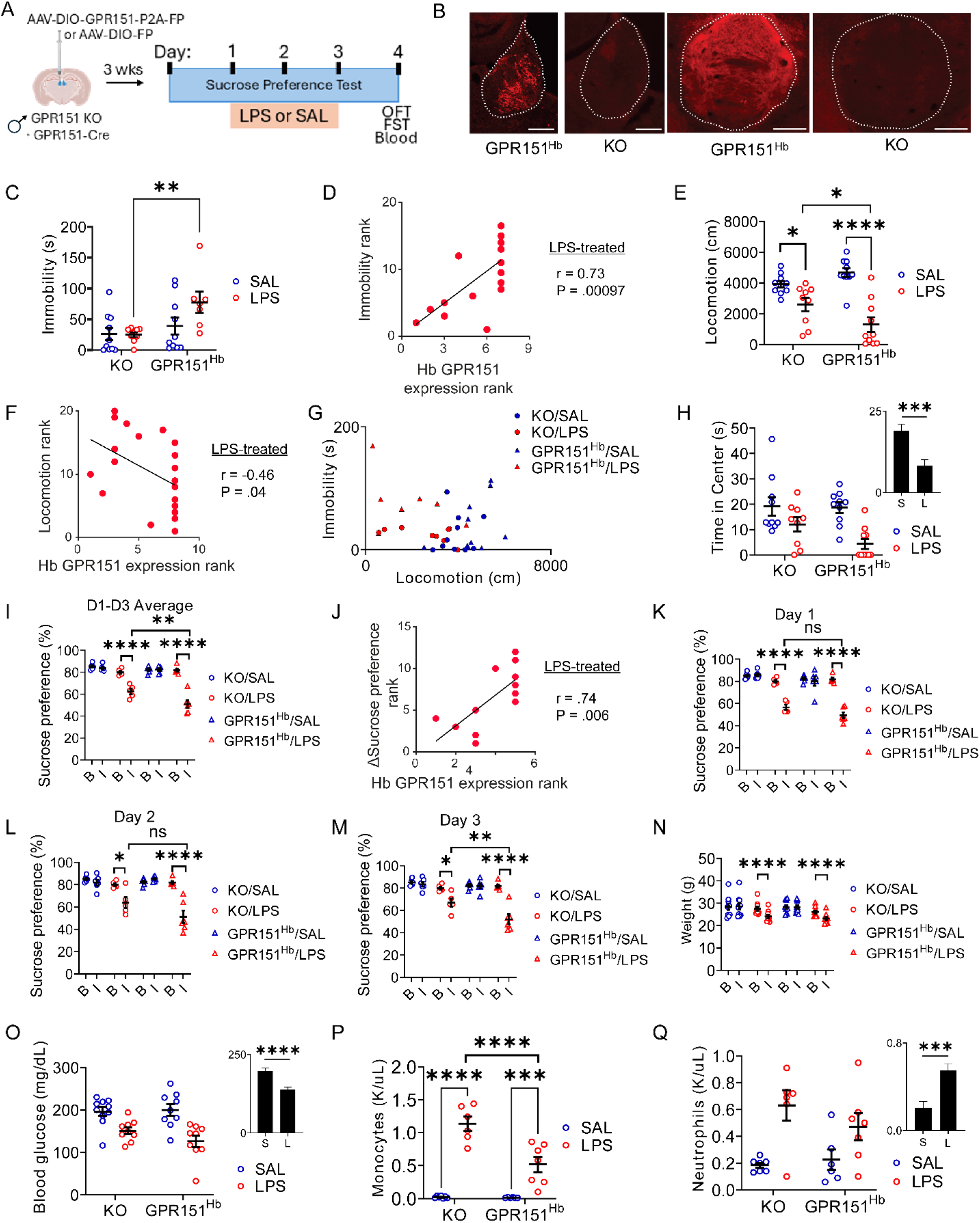
Behavioral and physiological responses to LPS in male *Gpr151* knockout mice with habenular re-expression of GPR151. **A**, Experimental methods. **B**, Example immunohistochemical labeling for GPR151 (red) in GPR151^Hb^ mice and mice injected with control AAV virus (‘KO’). Left, habenula is outlined. Right, midbrain at IPN. Scale, 0.2 mm. **C**, Immobility in the forced swim test. Planned comparisons, Mann-Whitney, KO/LPS vs GPR151^Hb^/LPS, P = .003, KO/Saline vs GPR151^Hb^/Saline, *P* = .3. *n* = 10 KO/Saline, 9 KO/LPS, 10 GPR151^Hb^/Saline, and 7 GPR151^Hb^/LPS mice. **D**, GPR151 expression in the habenula vs immobility in the forced swim test (shown as ranks; 1 is highest) in LPS-treated mice. *N* = 17 mice. **E**, Locomotion in the open field test. 2-way ANOVA, Virus x Injection, *P* = .008; KO/LPS vs GPR151^Hb^/LPS, *P* = .02; KO/Saline vs GPR151^Hb^/Saline, *P* = 0.15. *n* = 10 KO/Saline, 9 KO/LPS, 10 GPR151^Hb^/Saline, and 10 GPR151^Hb^/LPS mice. **F**, GPR151 expression in the habenula vs locomotion in the open field test (shown as ranks) in LPS-treated mice. *N* = 20 mice. **G**, Scatterplot of locomotion in the open field test and immobility in the forced swim test. *r* = -.21, *P* = .21. Mice as in C. **H**, Time in the center of the open field. Inset shows the Saline and LPS estimated marginal means, illustrating the main effect of Injection. 2-way ANOVA, Virus x Injection, P > .05. Mice as in E. **I**, Sucrose preference as in Fig. 2,3. 3-way RM ANOVA, Time x Virus x Injection, *P* = .01; KO/LPS vs GPR151^Hb^/LPS, *P* = .002. *n* = 6 KO/Saline, 5 KO/LPS, 6 GPR151^Hb^/Saline, and 6 GPR151^Hb^/LPS mice. **J**, GPR151 expression in the habenula vs change in sucrose preference (baseline – post-injections; shown as ranks) in LPS-treated mice. *N* = 12 mice. **K**, Sucrose preference for 24 hours after the first LPS injection. 3-way RM ANOVA, Time x Virus x Injection, *P* = .29. Note that ‘B’ is the same for I,K,L,M. **L**, Same as K, but after the second LPS injection. 3-way RM ANOVA, Time x Virus x Injection, *P* = .02; KO/LPS vs GPR151^Hb^/LPS, *P* = .052. **M**, Same as K, but after the third LPS injection. 3-way RM ANOVA, Time x Virus x Injection, *P* = .03; KO/LPS vs GPR151^Hb^/LPS, *P* = .003. **N**, Body weight, as in Fig. 2,3. KO/LPS/Injections vs GPR151^Hb^/LPS/Injections, *P* = .99. Mice as in E. **O**, Blood glucose, as in Fig. 2,3. Inset similar to H. 2-way ANOVA, Virus x Injection, P > .05. *n* = 10 KO/Saline, 9 KO/LPS, 9 GPR151^Hb^/Saline, and 9 GPR151^Hb^/LPS mice. **P**, Monocytes, as in Fig. 2,3. 2-way ANOVA, Virus x Injection, P = .001. *n* = 7 KO/Saline, 6 KO/LPS, 6 GPR151^Hb^/Saline, and 7 GPR151^Hb^/LPS mice. **Q**, Neutrophils, as in Fig. 2,3. Inset similar to H. 2-way ANOVA, Virus x Injection, P > .05. Mice as in P. Please see Supplementary Table 1 for more statistical information.

Despite the strong effects in males, the same viral microinfusions of GPR151-expressing AAV in the habenula of female *Gpr151* knockout/*Gpr151*-cre mice with the same LPS injection regimen (Fig. 5A,B; and performed concurrently with the experiments in males) did not increase inflammation-associated amotivation in any behavioral test (Fig. 5C-M), and GPR151 expression in LPS-treated females was not correlated with immobility in the forced swim test, locomotion, nor sucrose preference (Fig. 5D,F,J; Supplementary Fig. 3A-F), indicating that habenular re-expression of GPR151 was not sufficient to increase inflammation-associated amotivation in female mice under these conditions. However, LPS did cause slight changes in the open field test in females, as well as decreases in weight and blood glucose, and an increase in monocytes (Fig. 5E,H,N-P), indicating that LPS was still physiologically and behaviorally effective.

**Fig. 5.**
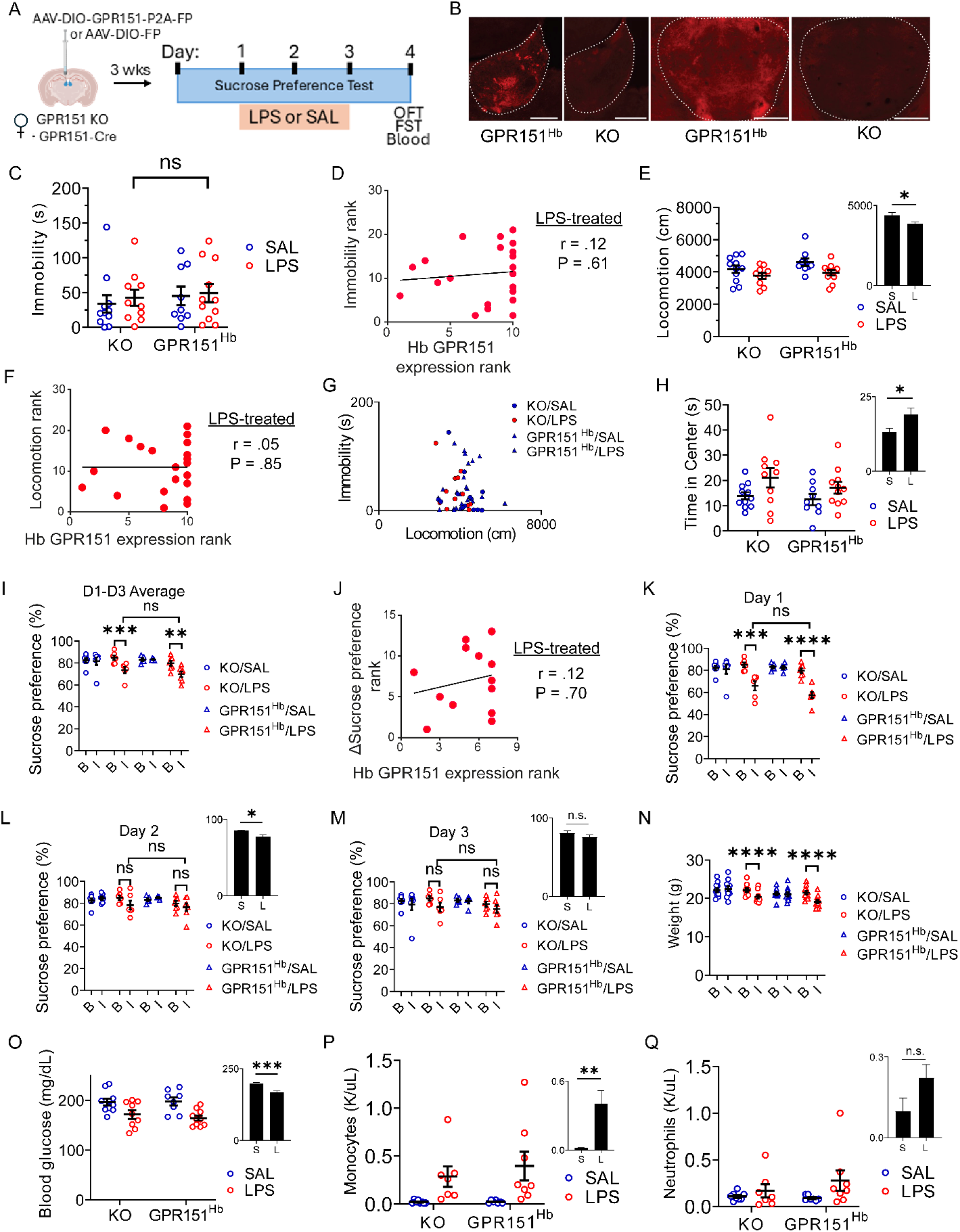
Behavioral and physiological responses to LPS in female *Gpr151* knockout mice with habenular re-expression of GPR151. **A**, Experimental methods, as in Fig. 4. **B**, Example GPR151 labeling, as in Fig. 4. **C**, Immobility in the forced swim test. Planned comparisons, Mann-Whitney, KO/LPS vs GPR151^Hb^/LPS, P > .05, KO/Saline vs GPR151^Hb^/Saline, *P* > .05. *n* = 11 KO/Saline, 10 KO/LPS, 9 GPR151^Hb^/Saline, and 11 GPR151^Hb^/LPS mice. **D**, GPR151 expression in the habenula vs immobility in the forced swim test (shown as ranks) in LPS-treated mice. *N* = 21 mice. **E**, Locomotion in the open field test. Inset shows the Saline and LPS estimated marginal means, illustrating the main effect of Injection. 2-way ANOVA, Virus x Injection, P > .05. Mice as in C. **F**, GPR151 expression in the habenula vs locomotion in the open field test (shown as ranks) in LPS-treated mice. *N* = 21 mice. **G**, Scatterplot of locomotion in the open field test and immobility in the forced swim test. *r* = −0.10, *P* = .47. Mice as in C. **H**, Time in the center of the open field, as in Fig. 2-4. Inset similar to E. 2-way ANOVA, Virus x Injection, P > .05. Mice as in C. **I**, Sucrose preference, as in Fig. 2-4. 3-way RM ANOVA, Virus x Injection x Time, P > .05. *n* = 7 KO/Saline, 6 KO/LPS, 5 GPR151^Hb^/Saline, and 7 GPR151^Hb^/LPS mice. **J**, GPR151 expression in the habenula vs change in sucrose preference (baseline – post-injections; shown as ranks) in LPS-treated mice. *N* = 13 mice. **K**, Sucrose preference for 24 hours after the first LPS injection. 3-way RM ANOVA, Virus x Injection x Time, P > .05. **L**, Sucrose preference for 24 hours after the second LPS injection. Inset similar to E. 3-way RM ANOVA, Virus x Injection x Time, P > .05. **M**, Sucrose preference for 24 hours after the third LPS injection. Inset similar to E. 3-way RM ANOVA, Virus x Injection x Time, P > .05. **N**, Body weight, as in Fig. 2-4. 3-way RM ANOVA, Virus x Injection x Time, P > .05. Mice as in C. **O**, Blood glucose, as in Fig. 2-4. Inset similar to E. 2-way ANOVA, Virus x Injection, P > .05. *n* = 10 KO/Saline, 9 KO/LPS, 8 GPR151^Hb^/Saline, and 10 GPR151^Hb^/LPS mice. **P**, Monocytes, as in Fig. 2-4. Inset similar to E. 2-way ANOVA, Virus x Injection, P > .05. *n* = 8 KO/Saline, 7 KO/LPS, 7 GPR151^Hb^/Saline, and 8 GPR151^Hb^/LPS mice. **Q**, Neutrophils, as in Fig. 2-4. Inset similar to E. 2-way ANOVA, Virus x Injection, P > .05. Mice as in P. Please see Supplementary Table 1 for more statistical information.

### LPS decreases *Gpr151* expression in the habenula

To determine if *Gpr151* expression is sensitive to LPS-induced inflammation, we repeated the three-day LPS treatment on a new group of wild-type mice and measured habenular *Gpr151* expression with qPCR (Fig. 6A). LPS reduced *Gpr151* expression in the habenula, without any effect of sex (Fig. 6B), indicating that *Gpr151* expression in the habenula is sensitive to inflammatory challenge in male and female mice.

**Fig. 6.**
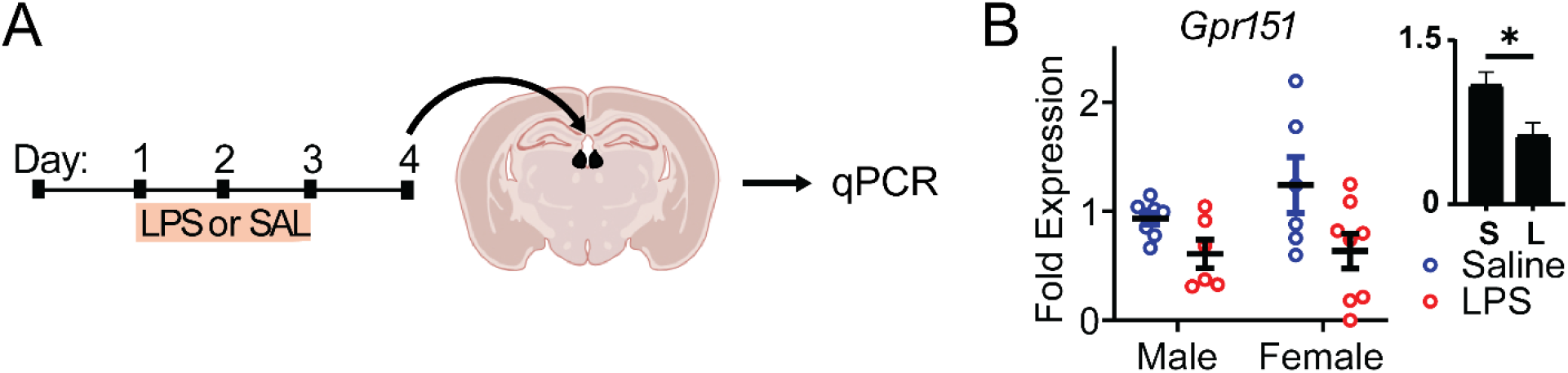
The effect of LPS on habenular *Gpr151* expression. **A**, Experimental methods. **B**, Fold expression of *Gpr151* relative to mean *Gpr151* expression in saline mice (males and females combined). 2-way ANOVA using a linear mixed effect model, Main effect of Injection, F(1,24) = 6.3, P = .02. *n* = 8 SAL/Male, 6 LPS/Male, 6 SAL/Female, 8 LPS/Female samples, 2 samples/mouse accounted for by linear mixed effect model. Sex x Injection, F(1,24) = .7, P = .42; Main effect of Sex, F(1,24) = 1.6, P = .21. Inset shows estimated marginal means with standard error to illustrate the main effect of Injection.

## Discussion

In this study, we find that male and female *Gpr151* knockout mice have reduced immobility in the forced swim test compared to wild-type mice after LPS treatment. This was the only behavioral measure for which there was an interaction between genotype (WT/KO) and injection (SAL/LPS), indicating that GPR151 regulates inflammation-associated changes in stress coping behavior—a behavior that is affected oppositely by chronic stress^80,81^, a risk factor for depressive disorders^82–88^, and antidepressants^89,90^. This effect could not be explained by differences in general motor behavior induced by LPS because there was no interaction between genotype and injection on locomotion in the open field test, nor a relationship between immobility in the forced swim test and locomotion in the open field test. The specificity of the effect in the forced swim test was further supported by the observations that knockout mice lost similar amounts of body weight after LPS, and male knockout mice had similar activation of the immune system as control mice given LPS, as indicated by blood monocyte and neutrophil levels.

Re-expression of GPR151 in the habenula of adult male *Gpr151* knockout/*Gpr151*-cre mice increased immobility in the forced swim test after LPS compared to control mice, indicating that GPR151 expression in adults can affect behavioral sensitivity to inflammation. Furthermore, the levels of habenular GPR151 expression correlated with the amount of immobility in LPS-treated, but not saline-treated, mice. Re-expression of GPR151 in the habenula also increased LPS-induced amotivation in the open field test, as assessed by locomotion (although again there was no relationship between immobility in the forced swim test and locomotion in the open field test), and increased LPS-induced anhedonia in the sucrose preference test. Both of these effects were also correlated with habenular GPR151 expression in LPS-treated mice. Interestingly, re-expression of habenular GPR151 did not affect sucrose preference on Day 1 of LPS injections, but did affect sucrose preference on subsequent days. This is consistent with a model wherein habenular GPR151 contributes to longer-lasting, inflammation-associated depression, but not acute, sickness responses to LPS, which may be mediated by other mechanisms^91–93^. Notably, GPR151 was virally re-expressed using a strong promoter (synapsin) on a knockout background that may have undergone compensation for the loss of GPR151. This could potentially explain the strong effect of GPR151 re-expression on LPS-induced changes in behavior relative to wild-type mice. Furthermore, we found that *Gpr151* expression is reduced by LPS in wild-type mice, which could also contribute to differences in behavior between GPR151^Hb^ and wild-type mice.

Unexpectedly, despite the strong effects of habenular re-expression of GPR151 in male mice treated with LPS, the same methods in female mice had no observable effects on motivated behavior. Female mice in our study were also less affected by LPS than male mice, as evidenced by less hypoactivity in the open field test, less anhedonic behavior in the sucrose preference test, and less of an increase in blood monocytes and neutrophils. Therefore, it is possible that our conditions were not optimal for measuring inflammation-associated amotivation in females. It is also possible that there was too much variability in the behavior of control female mice, perhaps exacerbated by surgery, and especially in the forced swim test, to measure differences between groups. Another possibility is that habenular GPR151 is involved in inflammation-associated amotivation in female mice, but is not sufficient to alter behavior in response to LPS because the receptor is also needed elsewhere. Consistent with this possibility, female knockout mice given LPS had reduced levels of blood monocytes and neutrophils relative to wild-type mice—an effect that was not restored by re-expression of GPR151 in the habenula. Finally, it is possible that there are sex differences in habenular circuitry^94^ that contribute to inflammation-associated amotivation or differences in the function of GPR151-expressing habenular neurons in male and female mice. Future studies examining sex differences in the role of GPR151-expressing habenular neurons in motivated behavior and the function of GPR151 outside the habenula, as well as loss-of-function experiments in the habenula and studies using optimized protocols for female mice are needed to test these possibilities.

Previous studies indicate that *Gpr151* is an injury- and inflammation-responsive gene in peripheral sensory pathways, with robust induction in sensory ganglia after neuropathic injury and in lung-innervating vagal sensory neurons after LPS-induced pulmonary inflammation^60–62,66,68–70^. Consistent with a role in the effects of inflammation, we found that LPS-treated female knockout mice had lower levels of blood monocytes and neutrophils than LPS-treated female wild-type mice, and re-expression of GPR151 in the habenula also, paradoxically, caused a strong reduction in blood monocyte levels in male mice. However, we found that LPS caused a decrease in *Gpr151* expression in the habenula, possibly due to homeostatic processes, unlike prior studies which showed an increase in *Gpr151* expression in sensory ganglia during inflammation^60–62,68^. Other studies indicate that the function of GPR151 in the habenula extends beyond inflammation. For example, GPR151 in MHb neurons affects behavioral responding for nicotine and synaptic transmission in the midbrain^79^. Because GPR151 has constitutive G-protein signaling^79^, it may be involved in many situations that engage GPR151-expressing habenular neurons. Thus, the therapeutic potential of GPR151 may be broader than inflammation-associated depression. Consistent with this, we observed differences in open field behavior in male *Gpr151* knockout mice that were independent of LPS. We also note that *GPR151* has been genetically linked to body mass index in humans^95^ and affects glucose tolerance in mice^96^, although we saw no differences in body weight nor blood glucose in knockout mice in our conditions, and the available evidence does not yet establish a definitive causal role for GPR151 in body mass index, as follow-up human knockout and mouse experiments yielded conflicting results^97^.

Our results support GPR151 as a potential therapeutic target for inflammation-associated depression and localize its effect on behavioral sensitivity to inflammation to the habenula in adult male mice. However, it remains to be determined which GPR151-expressing habenular neurons are involved. The distinction between the MHb and LHb is particularly salient given their distinct hodologies^98^; however, there may be specialized functions even among different GPR151-expressing groups of neurons within each structure^73,99^. Intriguingly, ketamine blocks LPS-induced, depression-related behavior in male mice^100^, and other studies indicate that the antidepressant effects of ketamine are initiated by ketamine’s block of NMDA receptors specifically in the LHb^38,43,101,102^. Therefore, it is possible that *Gpr151* knockout specifically in the LHb would have a similar antidepressant effect. Future studies will further refine our understanding of the neural circuitry involved in inflammation-associated depression and the mechanistic role of GPR151 in these circuits.

## Acknowledgments

We thank Gustavo Turecki and Annie Baccichet at the Douglas-Bell brain bank for human habenula tissue, Shigeyoshi Itohara and Chie Sano for Gpr151-cre mice, Shari Birnbaum for help with behavior, Ram Madabhushi and lab members for help with qPCR, Newaz Ahmed and Gena Konopka for help with single-nucleus RNA sequencing, and Robert Malinow for comments on the manuscript. We also thank Lenora Volk, Gena Konopka, Madhukar Trivedi, and Jeff Zigman for helpful discussion. Cartoons were made with Biorender. We used GPT-5.6 to collate statistics for the supplementary table, which we reviewed, and we take full responsibility for the content. The work was supported by funds from the NIMH (R01MH120131) and UT Southwestern Medical Center.

## Author contributions

L.R., L.Y., Y.S., H.A., F.J., S.T., A.G., and R.R. collected data. L.R., Y.L., L.Y., and S.S. analyzed data. L.R. and S.S. wrote the manuscript.

