## Supplementary Methods and Figures for "A habenula-enriched GPCR, GPR151, regulates behavioral sensitivity to inflammation"

[1]Department of Psychiatry, University of Texas Southwestern Medical School, Dallas, TX, USA; [2]Department of Neuroscience, University of Texas Southwestern Medical School, Dallas, TX, USA; [3]Neuroscience Graduate Program, University of Texas Southwestern Medical School, Dallas, TX, USA; [4]Peter O'Donnell Brain Institute, University of Texas Southwestern Medical School, Dallas, TX, USA; [5]University of Texas at Dallas, Dallas, TX, USA; [6]Florida Atlantic University, Boca Raton, FL, USA

Contents:

Supplementary Methods and Materials

Supplementary Figures 1-3

Supplementary Methods and Materials:

**Surgery**

To restore GPR151 expression in the habenula, *Gpr151* knockout/*Gpr151*-cre mice received bilateral injections of a Cre-dependent AAV vector (pAAV.hSyn.DIO.Gpr151.T2A.mCerulean; produced in-house; serotype 1, 6.3 x $10^{12}$ gc/mL and serotype 9, 1.7 x $10^{12}$ gc/mL) or a control vector (pAAV.hSyn.DIO.GFP; serotypes 1 and 9; Addgene plasmid #50457). For each injection, a 1:1 mixture of AAV1 and AAV9 serotypes was combined immediately prior to surgery and delivered stereotaxically (control viruses were diluted to a concentration of 5.0 x $10^{12}$). Coordinates for the lateral habenula were AP: −1.8 mm, ML: ±0.5 mm, DV: −2.7 mm; for the medial habenula: AP: −1.8 mm, ML: ±0.34 mm (5° angle), DV: −2.5 mm, relative to bregma. Viral injections were performed at a rate of 100 nL/min using a nanoinjector pump, with 300 nL delivered per site. Mice were given carprofen (5 mg/kg, s.c.) once per day on the day of surgery and the following three days for pain management.

**Histology and immunohistochemistry**

At the conclusion of behavioral testing, mice were anesthetized with isoflurane and sacrificed by decapitation. Brains were rapidly dissected, flash frozen in liquid nitrogen, and stored at −80 °C. For histological verification, frozen brains were later immersion-fixed in 4% paraformaldehyde for 48 h, cryoprotected in 30% sucrose until the tissue sank, embedded in OCT, and coronally sectioned at 40 µm using a cryostat. Sections were incubated with rabbit anti-GPR151 primary antibody (Sigma, 1:500) followed by Alexa Fluor 568-conjugated anti-rabbit secondary antibody (1:800). Fluorescence was visualized using epifluorescence microscopy. Mice were ranked according to the amount of GPR151 expression in the habenula, MHb, and LHb. Mice lacking detectable GPR151 expression in the habenula were excluded from categorical group comparisons but retained in analyses relating measured expression to behavior.

**Blood collection and analysis**

At sacrifice, ~30 µL of blood was collected into EDTA-coated microtubes during decapitation and processed within 2 h by the UT Southwestern ARC Diagnostic Laboratories to measure circulating monocyte and neutrophil levels. Blood glucose was measured at collection using a handheld glucometer (Accu-Chek, Roche Diagnostics).

**In situ hybridization**

In situ hybridization for Gpr151 mRNA was performed on 30-µm free-floating coronal brain sections containing the habenula using the RNAscope 2.5 HD Brown kit (Advanced Cell Diagnostics, Newark, CA, USA) following the manufacturer's instructions with minor adaptations for floating tissue. Mouse brains were fixed in 4% paraformaldehyde, cryoprotected in 30% sucrose, and sectioned coronally at 30 µm on a cryostat. Free-floating sections were stored in PBS containing 0.1% sodium azide until staining.

On the day of the assay, sections were washed in PBS and treated with hydrogen peroxide (Pretreatment 1) for 40–60 min at room temperature, followed by heat-mediated antigen retrieval (Pretreatment 2) for 10 min. Sections were mounted onto SuperFrost Plus slides, dried at 60°C, and treated with protease (Pretreatment 3) for 15 min at 40°C. Hybridization was performed using an RNAscope probe specific to mouse Gpr151 mRNA (ACD, Cat. # [Mm-Gpr151, Lot #17132A]), followed by sequential amplification steps (AMP 1–6) and chromogenic detection. Slides were counterstained with hematoxylin, blued in 0.02% ammonia water, dehydrated in graded ethanol, cleared in xylene, and coverslipped with DPX mounting medium. Negative controls were performed using a bacterial DapB probe, and positive controls used a probe against Ppib, following ACD guidelines.

For fluorescence in situ hybridization, we used 20-um coronal brain sections containing the habenula using the RNAscope HiPlex12 Reagent Kit v2 (Advanced Cell Diagnostics, Newark, CA, USA) following the manufacturer's instructions for fixed-frozen tissue with minor adaptations for vibratome-sectioned material. Wild-type female mice were transcardially perfused with 1X PBS followed by freshly prepared 4% paraformaldehyde (PFA) in 1X PBS. Brains were post-fixed overnight in 4% PFA at 4°C and sectioned the following day. Coronal sections of 20 µm thickness were cut on a vibratome and mounted onto SuperFrost Plus slides (Fisher Scientific, Cat. No. 12-550-15). Slides were baked at 60°C, then post-fixed in prechilled 4% PFA in 1X PBS for 15 min at 4°C, and dehydrated through a graded ethanol series (50%, 70%, 100%, 100%; 5 min each at room temperature). A hydrophobic barrier was drawn around each section using an ImmEdge pen (Vector Laboratories) and allowed to dry completely.

Target retrieval was performed by steaming slides in 1X RNAscope Target Retrieval Reagent at ≥99°C for 5 min, followed by rinsing in distilled water for 15 sec and transfer to 100% ethanol for 3 min, then drying at 60°C for 5 min. Sections were then incubated with RNAscope Protease III at 40°C for 30 min in a HybEZ II Hybridization Oven (ACD), followed by two washes in distilled water.

Hybridization was performed using the RNAscope HiPlex probe targeting mouse *Gpr151* mRNA assigned to detection tail T2 (RNAscope HiPlex Probe – Mm-Gpr151-T2, Cat. No. 317321-T2, Lot No. 26021A; ACD), diluted 1:50 in RNAscope HiPlex Probe Diluent and incubated at 40°C for 2 hours. Signal was amplified sequentially using RNAscope HiPlex Amp 1, Amp 2, and Amp 3 (30 min each at 40°C), followed by fluorophore detection using RNAscope HiPlex Fluoro T1–T4 v2 (15 min at 40°C), which labels the T2 channel with DyLight 550 (emission ~562 nm, orange). Sections were counterstained with DAPI (30 sec at room temperature) and coverslipped with ProLong Gold Antifade Mountant (Fisher Scientific, Cat. No. P36930). Negative and positive controls were performed using the RNAscope HiPlex12 Negative Control Probe and RNAscope HiPlex12 Positive Control Probe – Mm v2 (ACD), respectively, following the manufacturer's guidelines.

**qPCR**

Brain tissue was rapidly dissected and sectioned using a mouse brain matrix. The habenula was microdissected using RNase-free instruments, flash-frozen in liquid nitrogen, and stored at −80 °C until processing. Total RNA was extracted using the GeneAll® Hybrid-R™ RNA isolation kit according to the manufacturer's instructions. RNA concentration and purity were assessed using a NanoDrop spectrophotometer (Thermo Fisher Scientific). cDNA was synthesized from total RNA using Takara RNA to cDNA EcoDry™ Premix in 20 μL reaction volumes, following the manufacturer's protocol. Quantitative PCR (qPCR) was performed using SYBR Green Master Mix (Bio-Rad) in a 96-well plate format. Each 20 μL reaction contained 10 μL SYBR Green mix, 1.6 μL of 10 μM forward and reverse primer mix, 6.4 μL nuclease-free water, and 2 μL cDNA template. The primers for Gpr151 were GGCTGGTTCATCTGCAAGTCCT and GGTCACTTGCATACGCGAAGCA. Amplification was performed on a Bio-Rad real-time PCR system using standard cycling conditions according to the manufacturer's recommendations. Melt curve analysis was conducted to confirm amplification specificity. Relative gene expression was calculated using the 2^−ΔΔCt method, with Gapdh used as the endogenous control for normalization and further normalized to the mean expression levels in male and female controls injected with saline.

**Human tissue**

Postmortem human brain samples were provided by the Douglas–Bell Canada Brain Bank (www.douglasbrainbank.ca) under approved institutional protocols. Frozen human habenula tissue (30–50 mg) was dissected on dry ice, transferred to pre-chilled tubes, and stored at −80 °C until processing. For single-nucleus RNA sequencing, we included habenular tissue from a 54-year-old Caucasian male who died accidentally. The sample had a postmortem interval of 54.5 hours, a pH of 6.30, and a 9-hour refrigeration delay. No psychiatric diagnoses or substance use were reported, and toxicology at death was negative. For in situ hybridization, formaldehyde-fixed brain tissue was obtained through the UT Southwestern Willed Body Program. The donor was a 74 year-old female, with cause of death being metastatic lung cancer. The habenula was dissected, put in 30% sucrose, then O.C.T., frozen, and sliced (20-50 um) on a cryostat.

Before nuclei isolation for snRNA-sequencing, the presence of habenular tissue was verified by cutting a 40 μm section from the flash frozen tissue and mounting it directly on a glass slide. Sections were fixed in 4% paraformaldehyde (PFA) at room temperature and processed for slide-mounted immunohistochemistry (IHC). Tissue was incubated with rabbit anti-GPR151 primary antibody (Sigma SAB4500418-100UG) followed by goat anti-rabbit Alexa Fluor 488 secondary antibody (Fisher 111-545-003). Images were acquired to confirm GPR151-positive cells consistent with habenular morphology prior to homogenization. Because the human habenula can be difficult to microdissect accurately, this verification step ensured that downstream sequencing material originated from confirmed habenular tissue.

**Nuclei isolation for single-nucleus RNA sequencing**

Nuclei were isolated using a composite workflow adapted from Nagy et al., 2020 (Turecki lab) and the 10x Genomics Single Cell protocol. Briefly, confirmed habenula tissue was minced and gently dounced on ice in lysis buffer (10 mM Tris-HCl, 10 mM NaCl, 3 mM $MgCl_2$, 0.1% NP-40, 1 mM DTT, 40–100 U/mL RNase inhibitor). After 5 min on ice, the suspension was brought to 5 mL with lysis buffer, quenched with wash buffer (1× PBS, 2 % BSA, 0.25 % glycerol, RNase inhibitor), filtered through a 30 μm mesh, and pelleted (500 × g, 5 min, 4 °C).

When debris was evident, nuclei were further cleaned on a 29 % iodixanol (OptiPrep) cushion (12,000 × g, 30 min, 4 °C). Pellets were resuspended in wash buffer, briefly permeabilized in 0.1× permeabilization buffer (NP-40/Tween-20/digitonin; 2 min on ice), washed, and resuspended in 1× Nuclei Buffer supplemented with DTT and RNase inhibitor. Nuclei integrity was assessed by trypan-blue staining and light microscopy. Final concentrations were adjusted to ~1.5–2.0 × $10^3$ nuclei/µL, and approximately 3,000 nuclei were loaded per sample for library construction.

**Library preparation, sequencing, and primary processing**

Chromium Next GEM Single Cell Gene Expression (v1.1; 10x Genomics) libraries were prepared following manufacturer instructions and sequenced on an Illumina platform at the UT Southwestern Next Generation Sequencing Core. Base calls were demultiplexed using Cell Ranger ARC v2.0.1 (mkfastq), and libraries were processed with the GRCh38-2020-A-2.0.0 reference (cellranger-arc count --localcores 16 --localmem 64). The final dataset contained 11,094 nuclei after standard filtering. Jobs were executed on the UTSW BioHPC cluster via SLURM (1 node, 64–128 GB RAM). Filtered feature–barcode matrices from outs/filtered_feature_bc_matrix were used for downstream analyses.

**Quality control, normalization, and clustering**

Unless otherwise specified, default Cell Ranger ARC thresholds were applied. Additional filtering excluded nuclei with <200 detected genes or >3 % mitochondrial reads. Counts were normalized (10,000 counts per nucleus; log-transformed) and the ~2,000 most variable genes were used for principal-component analysis (PCA).

**Cluster annotation and habenular markers**

Filtered data were normalized using the NormalizeData function in Seurat. Highly variable genes were identified using the variance-stabilizing transformation method (FindVariableFeatures, selection.method = "vst") with 2,000 variable features. The data were then scaled using ScaleData, and principal component analysis (PCA) was performed using the variable features (RunPCA). A shared nearest neighbor graph was constructed using the first 20 principal components (FindNeighbors, dims = 1–20), and clusters were identified using the Louvain algorithm implemented in FindClusters with a resolution parameter of 0.2. UMAP dimensionality reduction was used for visualization (RunUMAP, dims = 1–20). Differentially expressed genes for each cluster were identified using the FindAllMarkers function with min.pct = 0.25 and logfc.threshold = 0.25.

To identify neuronal populations within the habenula dataset, expression of known neuronal markers was examined using FeaturePlot. Based on previously reported neuronal markers in habenular transcriptomic studies, cells expressing STMN2 and THY1 were annotated as neurons (Hashikawa et al., 2020[1]) and extracted for further analysis.

To distinguish lateral habenula (LHb) and medial habenula (MHb) neurons, expression of subtype-enriched markers was examined. PCDH10, previously reported as enriched in LHb neurons[1], was used as a marker for LHb neurons, whereas CHRNB4, identified as a marker gene for MHb subpopulations in a human habenula transcriptomic study[2], was used to identify MHb neurons. Cells were assigned to LHb or MHb neuronal populations based on enrichment of these marker genes and subsequently extracted for downstream analyses. The same analysis was run on data from our human sample and publicly available data from Yalcinbas et al., 2025[2].

**Statistical analysis**

All analyses were conducted using GraphPad Prism 10 (GraphPad Software, San Diego, CA, USA). Data are presented as mean ± SEM. Data were analyzed using two and three-way ANOVA as indicated, followed by post hoc multiple comparisons tests (Fisher's LSD for standard ANOVAs and Sidak's test for repeated-measures ANOVAs, unless otherwise stated). All tests were two-tailed, and statistical significance was defined as $p < 0.05$. Investigators remained blinded to genotype and treatment during behavioral scoring and data analysis. Insets are used to show statistical effects that cannot be shown in the main panels (e.g., main effects).

**Differential search analysis of Allen Brain Atlas**

Differential search of the Allen Brain Atlas in situ hybridization database was performed at https://mouse.brain-map.org/ using manual search for each brain region (N = 220 brain regions) with contrast structure = 'grey'. The most highly enriched GPCR was recorded along with its enrichment value for each brain region. Importantly, only GPCRs/regions with enrichment values greater than 50 were checked for artifacts (bubbles/blemishes) in the target brain region. If an artifact was found, the next most highly enriched GPCR and its enrichment value was substituted for that brain region. Information on image alignment and quantification can be found here (https://brain-map.org/support/documentation/api-for-mouse-brain-atlas).

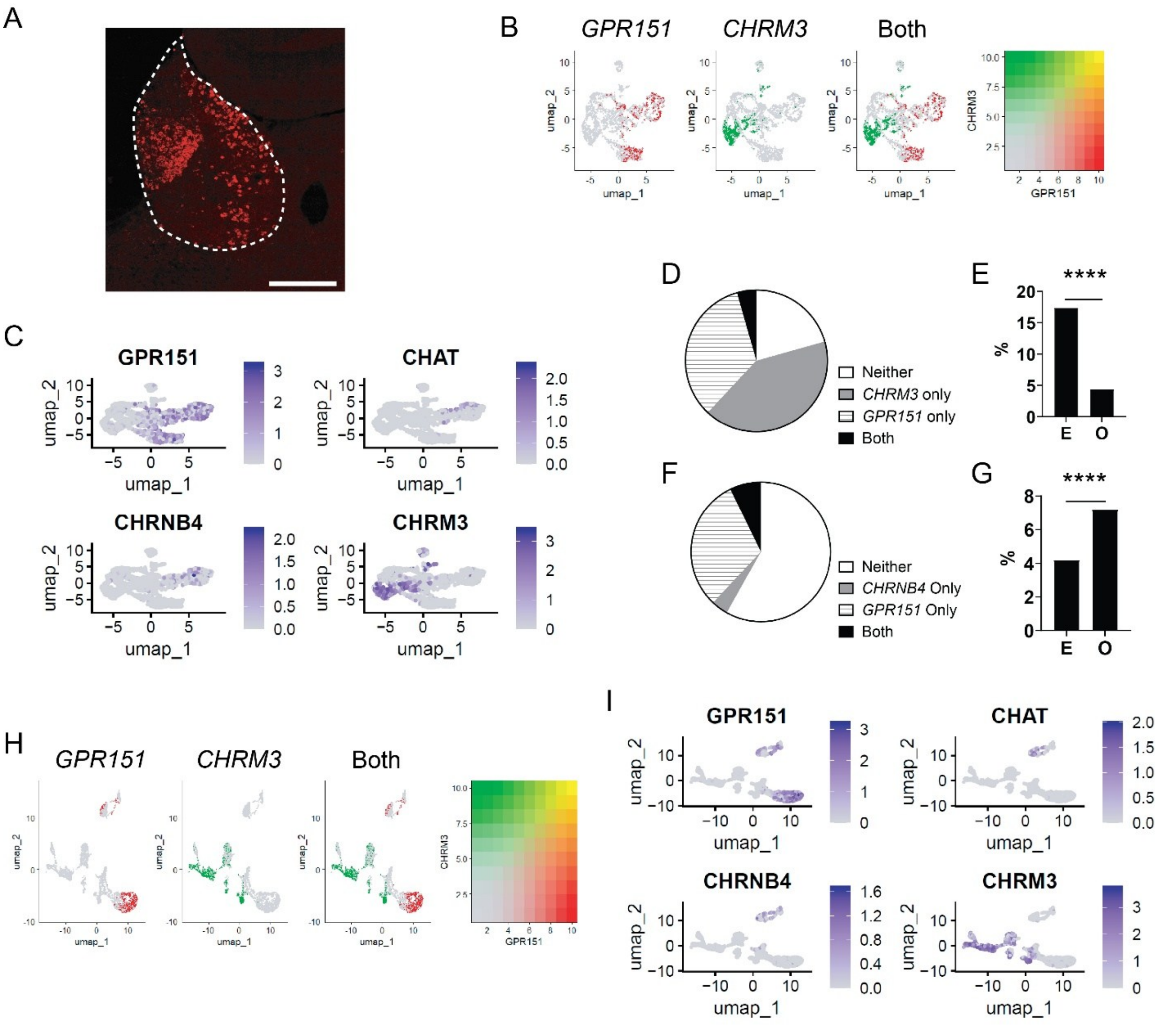


**Supplementary Figure 1. Data related to Figure 1. A**, Fluorescence in situ hybridization for *Gpr151* in a female mouse, showing expression in the habenula (dashed outline). Scale, 0.2 mm. **B**, UMAP plots of snRNA-seq expression in human habenula neurons showing expression of *GPR151* and *CHRM3*. **C**, UMAP plots, as in B, showing expression of *GPR151*, *CHRM3*, *CHRNB4*, and CHAT. **D**, Proportion of human habenula neurons from Yalcinbas et al., 2025[2] that express *GPR151* and/or *CHRM3*. $\chi 2 = 414.6$, $P = 4.9 \times 10^{-92}$, $N = 4781$ cells from 7 donors. **E**, 'E' denotes expected co-expression of *GPR151* and *CHRM3* based on the joint probability of their individual expression likelihoods. 'O' denotes observed co-expression. **F**, same as D but for *GPR151* and *CHRNB4*. $\chi 2 = 40.3$, $P = 2.2 \times 10^{-10}$, $N = 4781$ cells from 7 donors. **G**, same as E, but for *GPR151* and *CHRNB4*. **H**, UMAP plots, as in B, but using data from Yalcinbas et al., 2025[2]. N as in D. **I**, UMAP plots, as in C, but using data from Yalcinbas et al., 2025[2]. N as in D. ****, P < .0001.

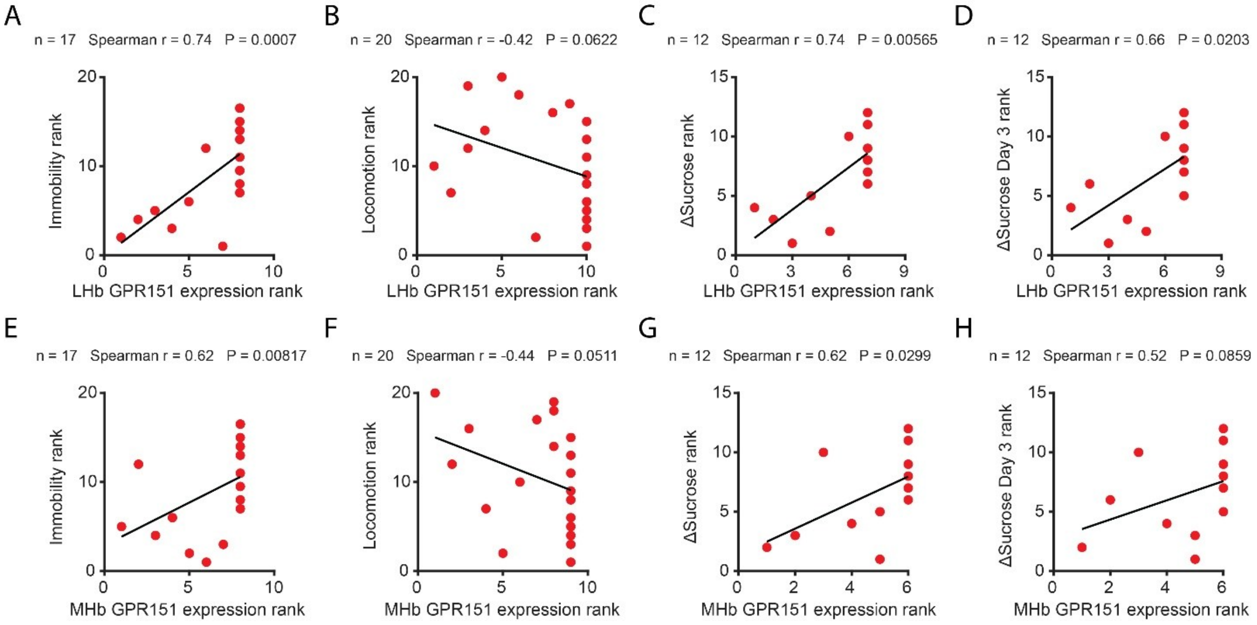


**Supplementary Figure 2. Data related to Figure 4.** Scatterplots of GPR151 expression in the LHb (A-D) and MHb (E-H) vs. behavior (shown as ranks) in male mice.

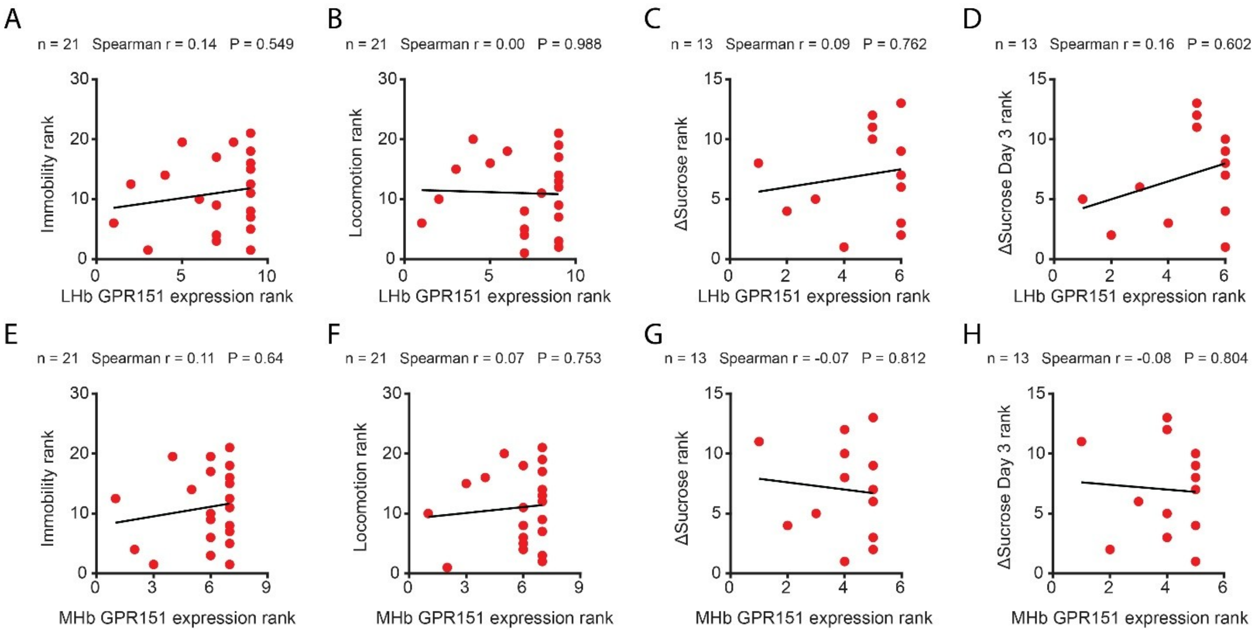


**Supplementary Figure 3. Data related to Figure 5.** Scatterplots of GPR151 expression in the LHb (A-D) and MHb (E-H) vs. behavior (shown as ranks) in female mice.

Supplementary Material References:

1 Hashikawa, Y. *et al*. Transcriptional and Spatial Resolution of Cell Types in the Mammalian Habenula. *Neuron* **106**, 743-758 e745 (2020). https://doi.org/10.1016/j.neuron.2020.03.011

2 Yalcinbas, E. A. *et al*. Transcriptomic Analysis of the Human Habenula in Schizophrenia. *Am J Psychiatry* **182**, 991-1006 (2025). https://doi.org/10.1176/appi.ajp.20240776
